# Jianpi Jiedu Recipe inhibits proliferation through reactive oxygen species-induced incomplete autophagy and reduces PD-L1 expression in colon cancer

**DOI:** 10.1101/2024.02.15.580471

**Authors:** Lingling Cheng, Liangfeng Xu, Hua Yuan, Qihao Zhao, Wei Yue, Shuang Ma, Xiaojing Wu, Dandan Gu, Yurong Sun, Haifeng Shi, Jianlin Xu

## Abstract

**Background:** Jianpi Jiedu Recipe has been used to treat digestive tract tumors in China since ancient times, and its reliability has been proven by clinical research. Currently, the specific biological mechanism of JPJDR in treating tumors is unclear.

**Methodology:** CCK-8 assay was used to detect cell viability. Clone formation assay and EdU assay were used to detect cell proliferation potential. DCFH-DA probe and JC-1 probe were used to detect total intracellular reactive oxygen species and mitochondrial membrane potential, respectively. Western blotting and immunofluorescence were used to detect protein expression level and subcellular localization of cells. RFP-GFP-LC3B reporter system was used to observe the type of autophagy in cells. Xenograft tumor model was used to study the therapeutic effect of JPJDR in vivo.

**Results:** JPJDR has a good inhibitory effect on various colorectal cancer cells and effectively reduces the proliferation ability of HT29 cells. After treatment with JPJDR, the amount of reactive oxygen species in HT29 cells increased significantly and the mitochondrial membrane potential decreased. JPJDR induced the accumulation of autophagosomes in HT29 cells and was shown to be incomplete autophagy. At the same time, JPJDR reduced the expression of PD-L1.Meanwhile, JPJDR can exert a good therapeutic effect in xenograft tumor mice.

**Conclusion:** JPJDR is a low-toxic and effective anti-tumor agent that can effectively treat colon cancer in vitro and in vivo. Its mechanism may be through inducing mitochondrial dysfunction and incomplete autophagy injury to inhibit the proliferation of colon cancer cells.

## 1. Introduction

Colorectal cancer (CRC) is one of the most commonly diagnosed cancers globally, posing a significant threat to global public health. According to data from the American Cancer Society, an estimated 153,020 new cases of colorectal cancer will be diagnosed in 2023, and 52,550 people will die from the disease[1]. The five-year survival rate for colorectal cancer is currently 58%-65%, which seems to be a comforting figure, as malignancies such as lung cancer, pancreatic cancer, and liver cancer currently have five-year survival rates of less than 20%[1,2]. However, this can only partially attributed to the development of medical care, as the five-year survival rate for colorectal cancer was 48%-51% in 1975, indicating that our treatment options for colorectal cancer remain limited[1]. Among patients diagnosed with metastatic colorectal cancer, fewer than 20% survive for more than 5 years. For unresectable metastatic colorectal cancer, Chemotherapy is the mainstay of therapy, as drugs can often diffuse to various parts of the body[3]. Currently, there is a lack of specific markers to identify high-risk patients who may benefit from adjuvant chemotherapy, and the efficacy of chemotherapy is often limited[4]. Therefore, there is an urgent need to find new drugs for the treatment of colorectal cancer.

China has used traditional Chinese medicine (TCM) to treat tumors for thousands of years and has demonstrated significant efficacy[5]. Additionally, natural compounds often have a richer structural and biological profile, making them an increasingly attractive resource for drug development[6]. Numerous studies have shown that the integration of TCM into the clinical treatment of tumors can significantly enhance treatment outcomes and reduce the occurrence of chemotherapy-related side effects[7,8]. Jianpi Jiedu Recipe (JPJDR) is a traditional Chinese medicine formula that is commonly used in China to treat gastrointestinal tumors. Recent studies have demonstrated that JPJDR can inhibit liver metastasis in colorectal cancer[9]. However, the specific mechanism by which JPJDR inhibits colorectal cancer remains unclear. It is urgently needed to elucidate its specific pharmacological effects to further guide its clinical application.

Reactive oxygen species (ROS) is a by-product of oxygen metabolism in living organisms, and the most important one is peroxide, which has been shown to maintain and activate some signaling transduction in tumor cells, promoting tumor progression at multiple stages[10]. Current research indicates that increasing the production of active oxygen can effectively kill tumor cells on the basis of their original high levels of active oxygen[11]. However, tumor cells experiencing oxidative stress can inactivate excessive active oxygen through autophagy and dispose of some damaged organelles and proteins to maintain internal environment stability[12]. Therefore, clinical trials have reported that blocking autophagy by chloroquine or hydroxychloroquine combined with chemotherapy drugs can produce better therapeutic effects[13]. Some studies have shown that blocking autophagy can increase the expression of PD-L1 in gastric cancer[14]. Typically, high expression of PD-L1 helps tumor cells evade immunity. Chemotherapy or antibodies that reduce PD-L1 expression have shown great potential as a treatment strategy and have been used in clinical trials[15]. Although PD-L1 expression does not significantly correlate with the prognosis of colorectal cancer[16], these results still suggest that blocking autophagy as a treatment strategy may hide some concerning risks. This study demonstrates that JPJDR plays a significant regulatory role in a series of cellular activities including active oxygen, mitochondrial membrane potential, autophagy, and PD-L1.

## 2. Materials and methods

### 2.1 Preparation of Jiangpi Jiedu decoction

The ingredients of JPJDR were purchased from Yancheng Traditional Chinese Medicine Hospital and identified by traditional Chinese medicine practitioner Lingling Cheng. The following are the traditional Chinese names, Latin names, and dosages of the ingredients:“Huangqi”Astragalus membranaceus (Fisch.) Bunge 30g, “Dangshen” *Codonopsis pilosula* (Franch.) Nannf. 30g, “Baizhu”Atractylodes Macrocephala Koidz. 15g, “Zhuling” Polyporus umbellatus (Pers.) Fr. 15g, “Yiyiren” *Coix lacryma-jobi L.var.mayuen(Roman.)Stapf* 15g, “Bayuezha” Fruit of Fiverleaf Akebia 15g, “Yeputaoteng” Vitis quinquangularis Rehd. 15g, “Hongteng” Sargentodoxa cuneata. 15g. All the herbs were soaked in water for 30 minutes and then boiled for 30 minutes. After that, they were continuously heated and evaporated until only 100ml of liquid remained. The herbal solution was filtered through gauze, then centrifuged at 14,000g for 1 hour. The supernatant was collected, filtered through a 0.22μm filter to sterilize, and stored at -80℃. The drug was then aliquoted and stored in this concentration of 1500 mg/ml.

### 2.2 Reagents and antibodies

The Cell Counting Kit-8 (Catalog number: C0039), EdU Cell Proliferation Kit with Alexa Fluor 488 (Catalog number: C0071S), Enhanced mitochondrial membrane potential assay kit with JC-1 (Catalog number: C2003S), and Reactive Oxygen Species Assay Kit (Catalog number: S0033M) was purchased from Beyotime Biotechnology(China). The following antibodies were all purchased from Cell Signaling Technology (USA): PCNA (Catalog number: 13110S), LC3B (Catalog number: 3868S), and P62 (Catalog number: 23214S). The following antibodies were all purchased from Proteintech Group (USA):GAPDH (Catalog number: 66248-1-Ig) and PD-L1 (Catalog number: 66248-1-Ig). The following reagents were all purchased from MedChemExpress (USA): Rapamycin (Catalog number: HY-10219) and Chloroquine (Catalog number: HY-17589A).

### 2.3 Cell Culture

HCT-116, HT-29, and SW480 cells were purchased from Wuhan Procell Life Science & Technology Co., Ltd (China). Cell culture was strictly carried out according to the manual provided by the company. HCT116 and HT29 cells were cultured in McCoy’s 5A (catalog number: PM150710), and SW480 was cultured in DMEM (catalog number: PM150210). The medium was supplemented with 10% fetal bovine serum (Gibico, 1009914C) without any antibiotics. Cells were placed in a constant temperature incubator with parameters set at 37℃, 5% CO2, and saturated humidity. The culture medium was changed every 2-3 days, and the cells were passaged once.

### 2.4 Cell vitality detection

The effect of JPJDR on cell viability was detected using the CCK-8 method. Cells were seeded in 96-well plates, with 10,000 cells per well, and incubated overnight. After the cells resumed their morphology the next day, the cells were treated with the specified concentration of drug for 24 or 48 hours. An appropriate amount of CCK-8 reagent was added to each well to make the working concentration 10%, and gently mixed. It was incubated in the incubator for 1 hour, and the optical density (OD) value of each well was measured using an enzyme microplate reader at a wavelength of 450 nm. The relative viability or inhibition rate of the cells was calculated based on the measured OD value.

### 2.5 Clone formation

The clone formation assay was used to detect the long-term effect of JPJDR on colon cancer cells. Briefly, 800 colon cancer cells were counted after treatment with a specified concentration of JPJDR for 24 hours and inoculated into a 6-well plate. Then, the cells were cultured in drug-free medium for 14 days, with the medium replaced every 3 days. After fixation with 3.7% paraformaldehyde, the cells were stained with 0.1% crystal violet, washed 5 times, and photographed.

### 2.6 Proliferation detection

The proliferation rate of cells was accurately detected using the EdU method. Colon cancer cells were seeded in a 6-well plate, and when the morphology recovered and the cells grew to 60% confluence, they were treated with a specified concentration of drug for 24 hours. EdU was added to the original culture medium at a two-fold concentration to make it a one-fold concentration. After culturing for another two hours, EdU was allowed to penetrate the replicating DNA. After fixing and permeabilizing the cells, a click reaction was performed to label EdU with Alexa Fluor 488. Finally, the cell nuclei were stained with DAPI and photographed.

### 2.7 Western blot

The relative protein abundance in cells was detected by Western blotting. After treating the cells with JPJDR for 24 hours, wash twice with PBS, and then lysate the cells at 4°C for 30 minutes using RIPA. Collect cell lysate and centrifuge at 14,000g for 30 minutes, and determine the protein concentration of each protein sample using the BCA method. Mix the same amount of protein lysate and loading buffer in the appropriate ratio, heat at 100°C for 5 minutes to denature the protein thoroughly. The prepared protein sample was subjected to SDS-PAGE gel electrophoresis, and then the protein was transferred to PVDF membrane using horizontal electrophoresis. 5% skimmed milk powder was used to block the gaps on the PVDF membrane, and it was washed with TBST for 3 times, each for 5 minutes. The primary antibody is usually incubated overnight at 4°C (usually for more than 14 hours). The secondary antibody is usually incubated at room temperature for 1 hour. Images were then acquired in the imager using chemiluminescence.

### 2.8 Immunofluorescence

The cells were seeded on special glass slides (the thickness must be less than 0.17mm) and cultured for 1-2 days to wait for the complete recovery of cell morphology. After 24 hours of treatment with JPJDR or control, the cells were fixed with 3.7% paraformaldehyde and then permeabilized with 0.1% Triton X-100. The blocking process was carried out in 5% skimmed milk and incubated at room temperature for 2 hours. According to the instructions of the primary antibody, the primary antibody was diluted with the appropriate dilution buffer, and the blocking solution was aspirated with blotting paper. Each glass slide was dropped with the diluted primary antibody and placed in a humid box, and incubated overnight at 4℃. After incubating the primary antibody, the cells were washed 3 times in PBST solution for 5 minutes each time. After washing, the secondary antibody labeled with fluorescence was added. According to the instructions of the secondary antibody, it was diluted with the appropriate dilution buffer and added to the glass slide for incubation. After incubating the secondary antibody, the cells were washed again and then observed under a fluorescence microscope. Before observation, the glass slide was dropped with a fluorescence quenching agent, and then sealed.

### 2.9 ROS detection

The determination of total intracellular ROS level was carried out by DCFH-DA probe. DCFH-DA probe itself has no fluorescence and can freely cross the cell membrane. After entering the cell, DCFH-DA probe is hydrolyzed into DCFH, which cannot freely cross the cell membrane, thus enriching the probe in the cells. The intracellular ROS oxidizes DCFH to DCF, and DCF can be excited to emit bright green fluorescence. The specific operation process is as follows: cells were cultured in 6-well plates, and when the cells recovered their morphology and reached about 60% confluence, they were intervened with drugs. After 24 hours, DCFH-DA probe was incubated with the cells for 30 minutes, washed with PBS once, and then collected cells are immediately detected on a flow cytometer. Due to the short lifespan and rapid change of ROS, to ensure the accuracy of the results, the total detection time should not exceed 1 hour from the time when the cells were digested and suspended.

### 2.10 Mitochondrial membrane potential detection

The JC-1 probe was used to detect mitochondrial membrane potential depolarization levels. When the mitochondrial membrane potential is at a high level of polarization, JC-1 accumulates in the mitochondrial matrix and emits red fluorescence. When the mitochondrial membrane potential is at a low level of polarization, JC-1 probe is scattered in the cytoplasm and emits green fluorescence. By comparing the ratio of red/green fluorescence intensity, it can be used to measure mitochondrial depolarization levels. Cells were cultured in 6-well plates, and when the cells recovered their morphology and reached about 60% confluence, they were treated with different concentrations of JPJDR or control conditions. After 24 hours, the culture medium was removed and fresh medium containing the JC-1 probe was added, and the cells were incubated for 40 minutes. Then, the cells were gently washed 3 times and imaged immediately using a fluorescence microscope.

### 2.11 Animal experiment

Thirty SPF-grade BALB/c nude female mice, 4-5 weeks old, were purchased from Changzhou Calvin Experimental Animal Co., Ltd (China). After 4 days of adjustable feeding, a volume of 200 μl PBS containing 1×10^7^ cells was injected subcutaneously into the mice. About 16 days after injection, 18 mice with tumors that had reached a volume of 100 mm^3^ were selected. The mice were randomly divided into 3 groups. The mice in the JPJDR group were administered a dose of 22g/kg (calculated based on the patient’s dosage), the mice in the positive control group were injected intraperitoneally with 5-FU (50mg/kg) every 3 days, and the model group mice were given an equal volume of water (290μl). On the 14th day, the mice were anesthetized, and the tumors were separated and fixed. During the experiment, all animals were kept in a SPF-grade barrier environment with a temperature of 22°C, relative humidity of 60%, and a 12-hour light cycle.

### 2.12 Immunohistochemistry

Briefly, the paraffin-embedded tissue is cut into a thickness of 4μm, adsorbed on a glass slide, heated at 72℃ for 2 hours. Subsequently, the tissue is gradually dewaxed in ethanol, and finally soaked in water for 5 minutes. The sections are heated by microwave for 3 minutes to repair the antigen, and 3% H_2_O_2_ is used to block endogenous peroxidase. The primary antibody is incubated overnight at 4℃, and the secondary antibody is incubated at room temperature for half an hour. The protein is quantitatively expressed by DAB.

### 2.13 Statistical analysis

All experiments were performed at least three times. Statistical analysis was conducted by GraphPad Prism software. Differences between three (or more) groups were analyzed using analysis of variance (ANOVA). Significance was designated as follows: *, P <0.05; **, P < 0.01; ***, P < 0.001.

## 3. Results

### 3.1 JPJDR inhibited CRC activity and proliferative potential

To preliminarily test the inhibitory effect of JPJDR on CRC, CCK-8 assay was used to detect the activity of the cells. The activity of JPJDR-treated HCT116,SW480, and HT29 cells was reduced in a dose-dependent manner compared to the control group, with IC50s of 0.77 mg/ml, 0.68 mg/ml, and 0.66 mg/ml, respectively (fig1A). Since HT29 cells were more sensitive to the drug treatment and had a good dose-inhibition curve fit, we chose HT29 for subsequent studies. JPJDR significantly reduced the clone formation ability of HT29 cells compared to the control group, indicating that their proliferative potential was down-regulated (fig1B). We next examined the proliferative capacity of HT29 using the EdU assay, which is a more sensitive way to detect proliferation. It showed that JPJDR significantly reduced the proliferative capacity of HT29 cells in a dose-dependent manner(fig2A,B). Meanwhile JPJDR substantially reduced the protein abundance of PCNA (fig2C), a proliferation marker for colorectal cancer. These results suggest that JPJDR has a favorable inhibitory effect on CRC.

**Fig.1.**
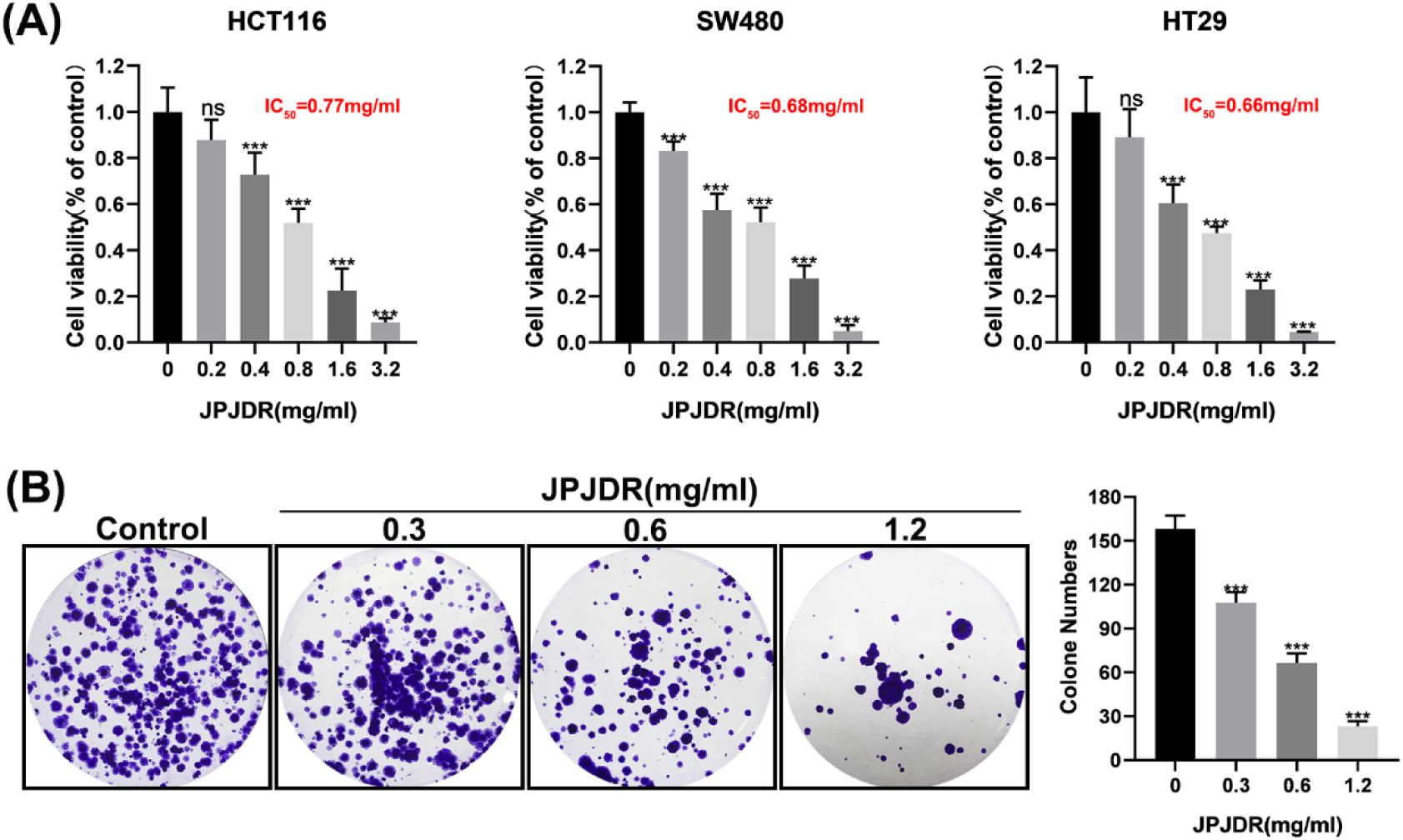
(A)Cell viability of HCT116, SW480, and HT29 after 24 hours of JPJDR treatment. (B)Representative images of HT-29 cell colony formation.

**Fig.2.**
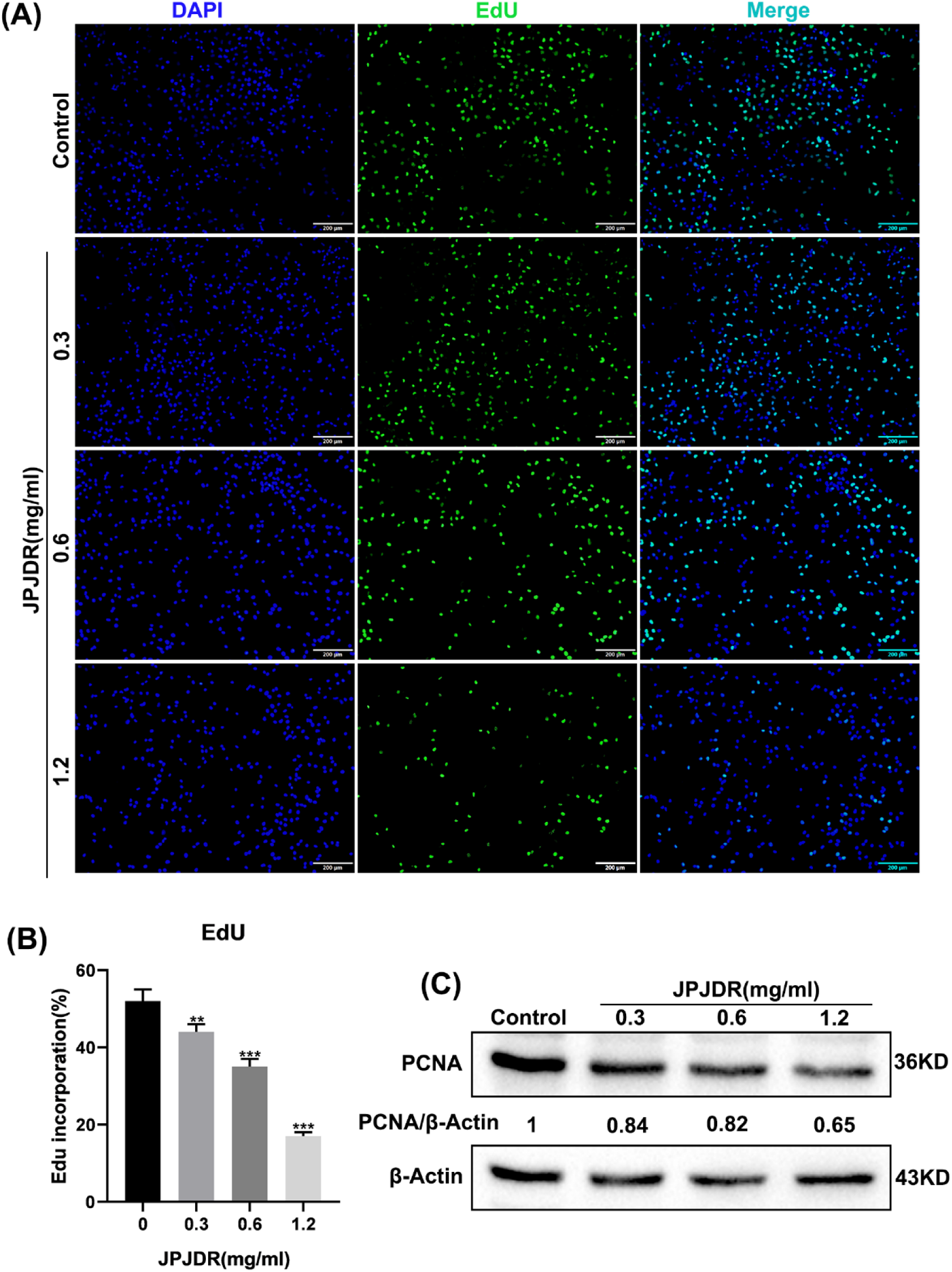
(A-B)EdU fluorescence image and statistical results of HT-29 cells. Scale bar 200 μm. (C)Immunoblotting image of proliferation marker PCNA. GAPDH was used as an internal reference.

### 3.2 JPJDR induced ROS accumulation and depolarized mitochondrial membrane potential in CRC cells

Subsequent studies found that JPJDR significantly increased reactive oxygen species levels in HT29 cells. Using the control group as the standard, thresholds for positive cell populations were set. Different concentrations of JPJDR increased the ROS-positive cell population from 12.1% in the control group to 23.0%, 37.5%, and 49.1%, respectively (fig3A,B). Studies have shown that ROS can disrupt mitochondrial membrane potential and impair mitochondrial function[17,18], and we subsequently examined the effect of JPJDR on mitochondrial membrane potential in HT29 cells using the JC-1 probe. In the control group, the JC-1 probe aggregated in the mitochondria and emitted red fluorescence, whereas after JPJDR treatment, the JC-1 probe was difficult to aggregate in the mitochondria and dispersed in the cytoplasm or outside of the mitochondria in the form of monomers which emitted green fluorescence (fig3C), suggesting that the mitochondrial membrane potential had been significantly depolarized.

**Fig.3.**
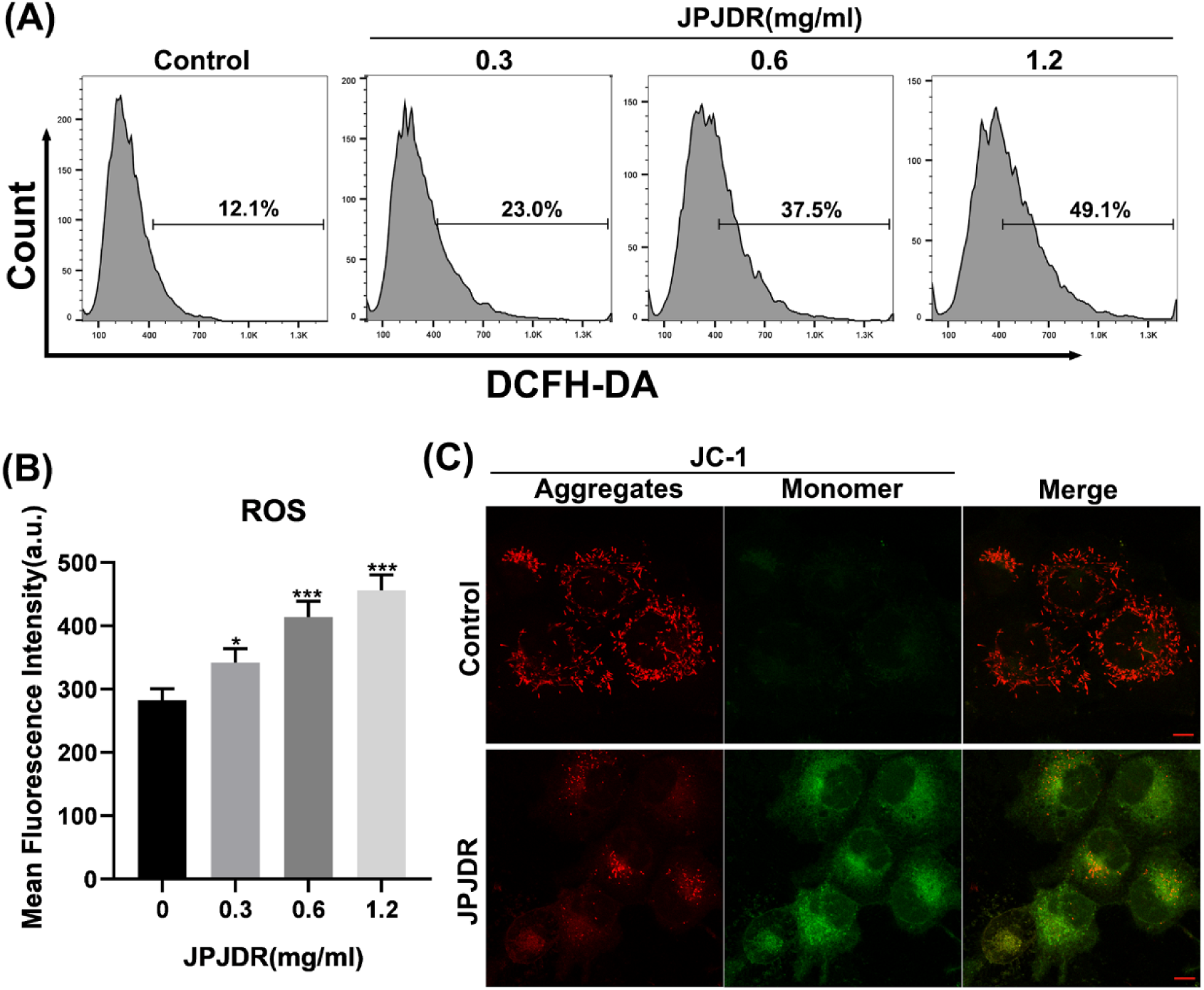
(A-B)The total reactive oxygen species level in HT29 cells was detected by the DCFH-DA probe in combination with flow cytometry. (C)Confocal microscopy imaging of HT29 cells incubated with the JC-1 probe. Scale bar 10 μm.

### 3.3 JPJDR induces incomplete autophagy in CRC cells

One key study has shown that depolarization of mitochondrial membrane potential strongly induces autophagy activation[19], while ROS has also been shown to be a powerful autophagy activator[20]. Therefore, we next examined the expression and subcellular distribution of LC3B protein, a specific marker for autophagic vesicles. When in the control state, LC3B protein was dispersed in the cytoplasm and nucleus at very low abundance.The protein abundance of LC3B in HT29 cells treated with JPJDR increased dramatically and aggregated into puncta in the cytoplasm (fig4A,B), a hallmark of autophagic vesicle formation.

**Fig.4.**
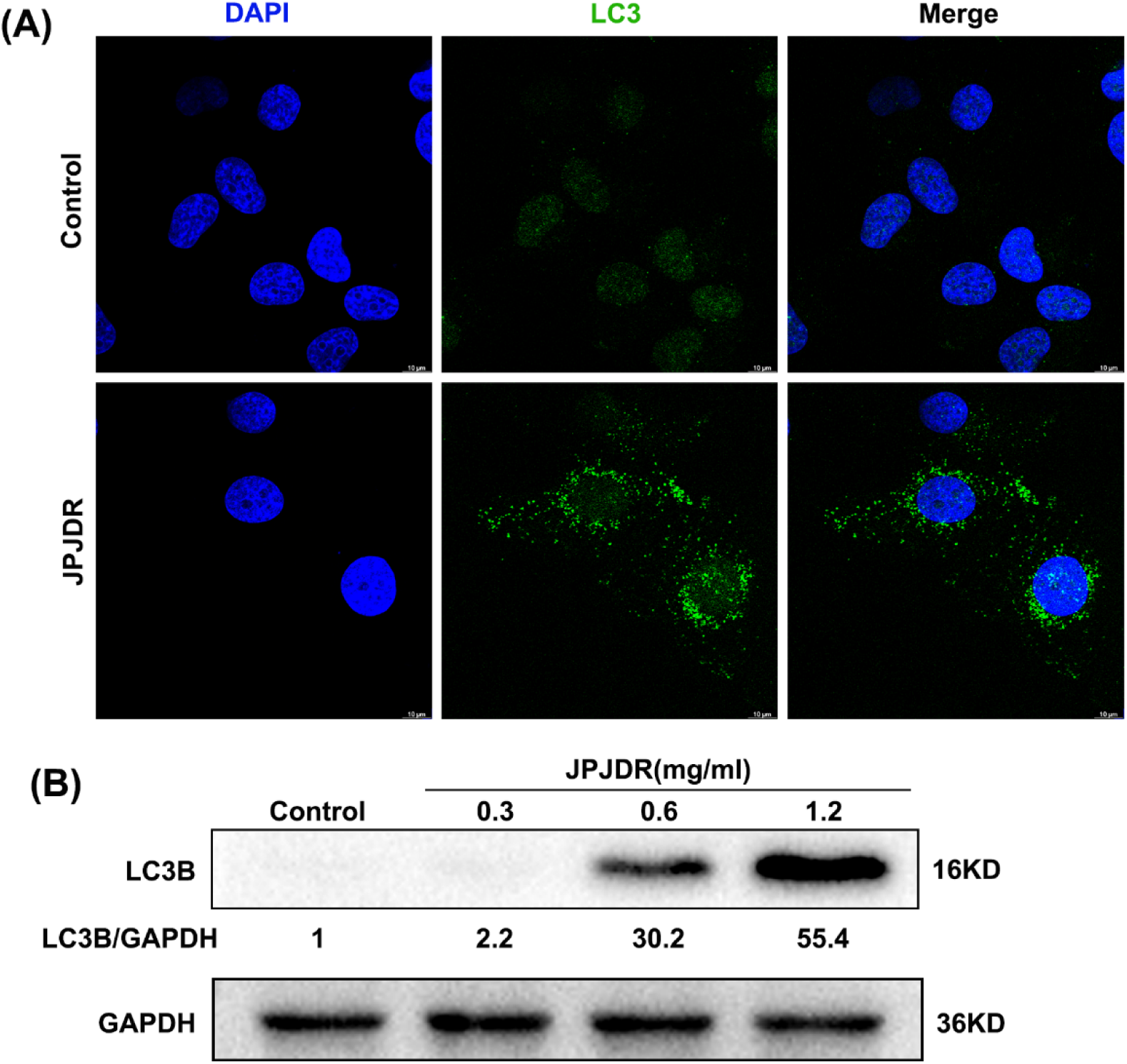
(A) Immunofluorescence images of LC3B protein in HT29 cells. Scale bar 10 μm. (B)The western blot image of LC3B protein in HT29 cells. GAPDH was used as an internal reference.

To investigate whether JPJDR increased autophagic flux, we reverse transcribed RFP-GFP-LC3B into HT29 cells. Rapamycin, a classical autophagy activator, induced an enhancement of autophagic flux through mTOR. Rapamycin-treated HT29 cells had a large number of red autophagic dots but only a very small number of green ones, indicating that the majority of autophagosomes are formed and have completed fusion with lysosomes. In contrast, red autophagic dots and green autophagic dots were co-expressed in JPJDR-treated HT29 cells, indicating that autophagosomes failed to fuse with lysosomes (fig5a). These results suggest that JPJDR blocked the autophagic flow in HT29 cells. To further elucidate whether JPJDR simply blocked autophagic flow or activated incomplete autophagy, we examined the abundance of p62 proteins. Co-upregulation of protein abundance of LC3B and P62 is a hallmark of autophagic flow blockade[21]. Chloroquine (CQ), a late autophagy blocker [22], significantly increased the protein abundance of LC3B and P62 (fig5b). In contrast, JPJDR more significantly induced upregulation of protein abundance of LC3B and P62. These results suggest that JPJDR induced incomplete autophagy in CRC.

**Fig.5.**
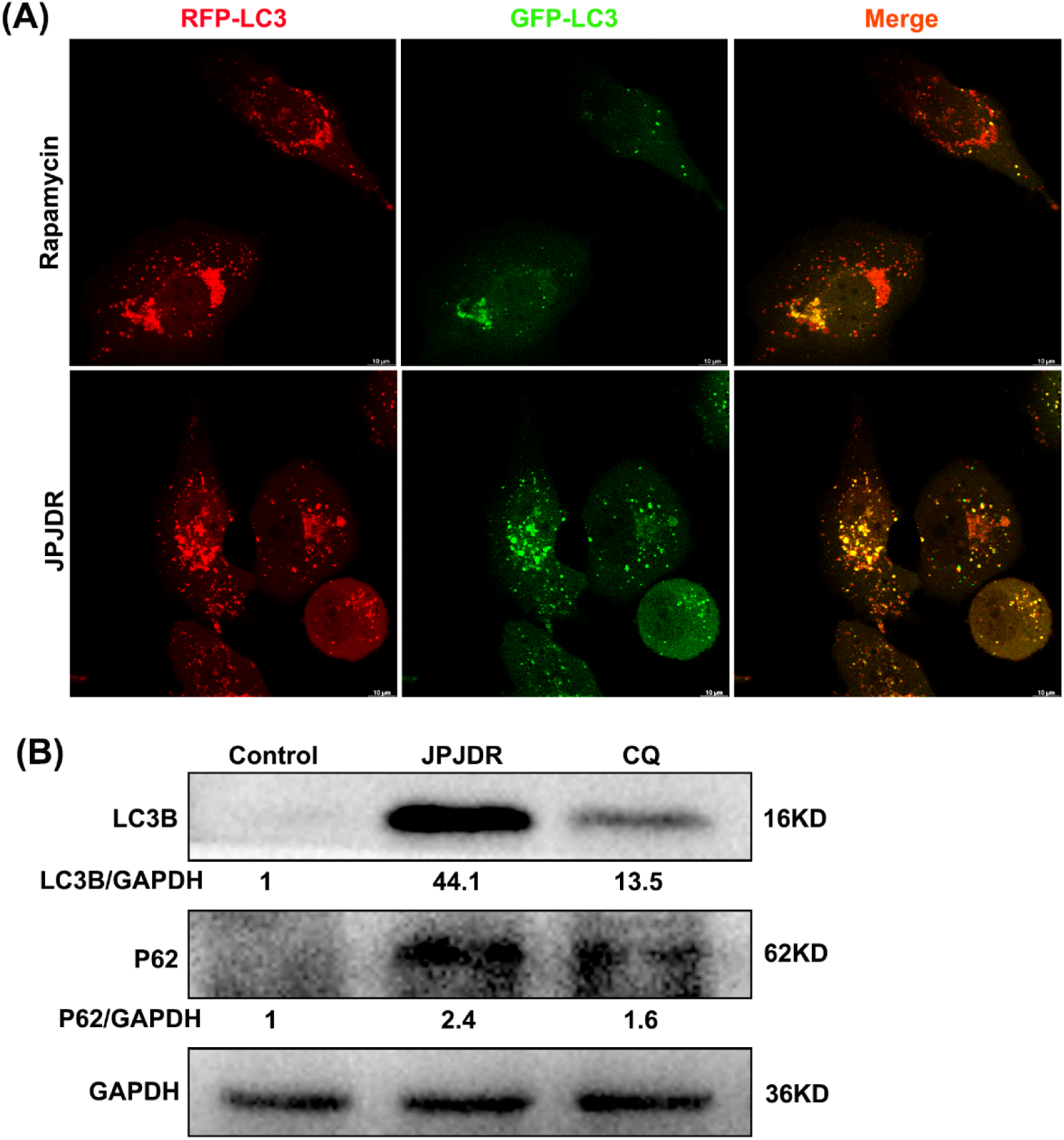
(A)Live cell imaging of HT29 cells expressing the RFP-GFP-LC3B sequence, captured by a confocal microscope. Scale bar 10 μm. (B)Immunoblotting images of LC3B and P62 proteins. GAPDH was used as an internal reference.

### 3.4 JPJDR reduced the expression of PD-L1 in CRC

The immunosuppression and immune escape of malignant tumors are among its ten characteristics. Programmed cell death ligand 1 (PD-L1) is an important means for cancer cells to escape immunity and is considered a potential target[23]. One key study showed that blocking autophagy increases PD-L1 expression in gastric cancer cells [14], so we focused on the effect of JPJDR on PD-L1 expression in CRC. After treatment with JPJDR, almost no PD-L1 expression was detected in HT29 cells, while more PD-L1 was detected in Chloroquine(CQ)-treated HT29 cells (fig6a). To more accurately detect PD-L1 protein abundance, we performed Western blot analysis. The results showed that JPJDR downregulated PD-L1 protein abundance in HT29 cells in a dose-dependent manner (fig6B). These results indicate that JPJDR can downregulate PD-L1 expression in CRC without upregulating PD-L1 due to autophagy.

**Fig.6.**
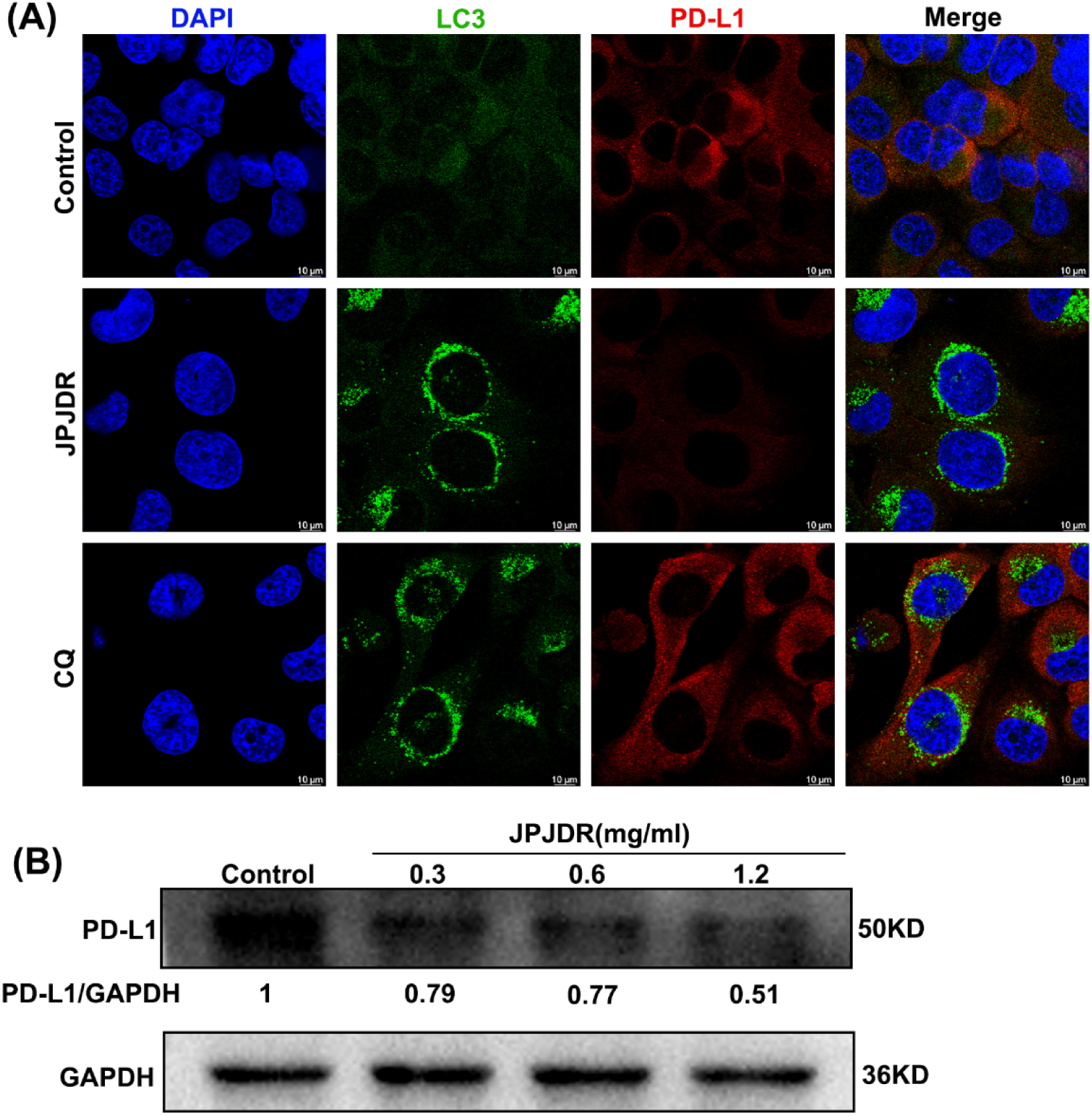
(A)Immunofluorescence images of LC3B(green) and PD-L1(red). Scale bar 10 μm. (B)Immunoblotting image of PD-L1. GAPDH was used as an internal reference.

**Fig.7.**
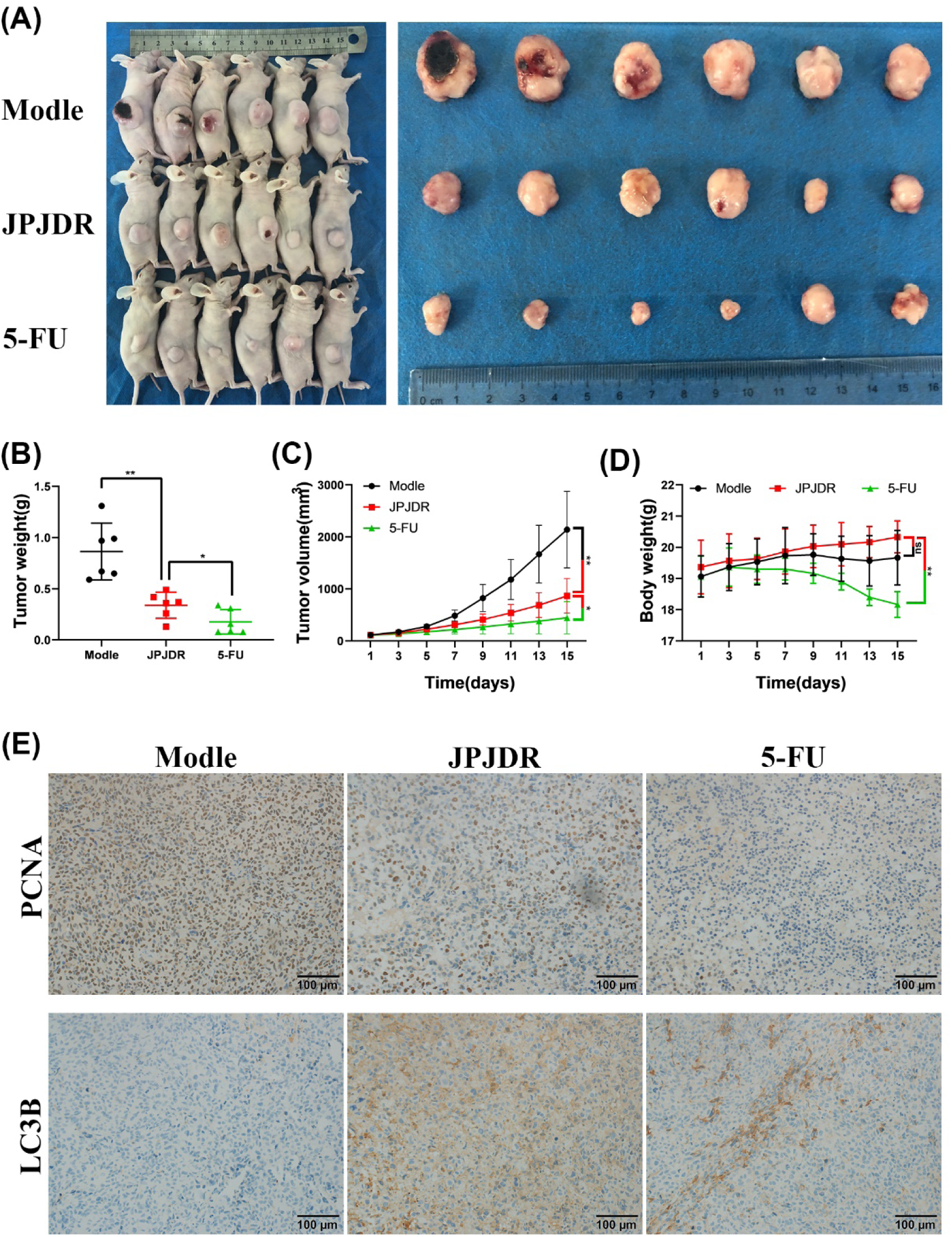
(A) Images of the whole mouse and isolated tumor. (B-D) Statistical analysis of tumor volume, weight, and mouse weight. (E) Immunohistochemical images of PCNA and LC3B proteins in tumor tissue. Scale bar 100 μm.

### 3.5 JPJDR inhibited colon cancer growth in vivo

Next, we constructed a xenograft tumor model of HT-29 cells to investigate whether JPJDR produces these effects in vivo. Compared with the control group, in mice treated with JPJDR for 14 days, the tumor volume was reduced from 2140.3±735.6 mm^3^ to 867.2±332.6 mm^3^, and the tumor weight was reduced from 0.86±0.28 g to 0.34±0.13 g, both of which were significant. The tumor-suppressing efficacy of JPJDR differed significantly from that of the positive drug 5-FU, with the volume and weight of the tumor after 14 days of 5-FU treatment being 446.3±0.28 g and 446.3±0.13 g, respectively. However, the weight of mice after 14 days of 5-FU treatment (18.2±0.4 g) was significantly lower than that of the JPJDR group (20.3±0.5 g).

Mouse tumors were isolated and subjected to immunohistochemical analysis. Compared with the control group, the positivity rate of tumor proliferation marker PCNA was significantly lower in the JPJDR group. Meanwhile, the accumulation of LC3B protein in tumor tissues was observed. These results preliminarily verified the consistency of JPJDR in the treatment of colon cancer in vitro and in vivo.

## 4. Discussions

Although in the past decades, the popularization of risk factors (such as smoking, increased use of aspirin, consumption of red meat and processed meat, obesity and diabetes) and the application of colonoscopic polypectomy have reduced the incidence rate and mortality, nearly 50% of patients after surgical resection have the risk of recurrence and metastasis leading to death within five years after diagnosis[3]. Due to the frequent occurrence of drug resistance events and patients’ intolerance to drugs, many colorectal cancer patients eventually face the dilemma of having no drugs available[11]. Therefore, it is urgent to find new drugs and strategies for the treatment of colorectal cancer.

Due to the high level of sugar metabolism, even with the presence of Warburg effect, the level of reactive oxygen species (ROS) remains high in tumor cells[24]. Previous research has shown that ROS can promote the occurrence and development of cancer, such as maintaining the activity of PI3K signaling[25]. Until 2011, one key study showed that ROS can relieve the progress of some cancers and found that the redox state in cells is an important factor determining tumor formation[26]. Now the mainstream view is that continuously increasing the generation of ROS in tumor cells is a preferred strategy to eliminate tumor cells[27,28], because the change of metabolic characteristics maintains a very dangerous high level of ROS in tumor cells. Mitochondria are the main site of ROS production, especially complex I. The increase of ROS can lead to the opening of mitochondrial membrane permeability transport pores, which directly leads to the reduction of mitochondrial membrane potential and induces apoptosis[29]. Long-term clinical applications show that JPJDR is a very low toxic compound preparation. This study found that JPJDR can increase the level of ROS in CRC cells and significantly reduce mitochondrial membrane potential, which may explain its effectiveness and low toxicity in treating cancer. Animal experiments have also shown that JPJDR did not significantly reduce the weight of the mice, unlike traditional chemotherapy drugs. On the contrary, 5-FU often induces digestive tract damage in clinical treatment, while JPJDR has been traditionally used to promote appetite and digestion.

Autophagy is a highly conserved cytoplasmic process that allows cells to "eat" a portion of themselves in order to counteract external environmental stress[22]. There are many inducers of autophagy, including energy deficiency, growth factor deficiency, accumulation of reactive oxygen species, microbial infection, organelle damage, and misfolded proteins[30]. Typically, the role of autophagy in cancer cells can be bivalent: cancer cells can relieve energy stress and abnormal proteins and organelles caused by drugs through autophagy, which is beneficial to their survival and drug resistance. However, if the autophagy process is interrupted, autophagosomes engulf various cell contents and accumulate in the cytoplasm without being digested, which can exacerbate damage and ultimately cause cell death, known as autophagic death[31]. Additionally, clinical applications have demonstrated that blocking autophagy is a good strategy for treating cancer[32]. Currently, there are a large number of compounds that have been shown to regulate autophagy, but their targets are relatively broad, which limits their clinical applications. JPJDR has long been used to treat gastrointestinal cancer, and its effectiveness and safety have been widely recognized. This study demonstrates that JPJDR is an incomplete autophagy activator. Compared with the simple autophagy late-phase inhibitor chloroquine, JPJDR causes more accumulation of LC3B Ⅱ in the cytoplasm. This study provides more evidence and application scenarios for the clinical application of JPJDR. Next, we will investigate what happens when drugs that are hindered by autophagy are combined with JPJDR.

Currently, antibodies or inhibitors against PD-1/PD-L1 are being tested in more than 1000 clinical trials, and some of them have been approved for cancer treatment [23]. Although one large clinical study in 2012 showed that PD-1 antibody treatment had no significant effect on colon cancer patients [16], a large number of studies have shown that PD-L1 blocking or ablation has good application potential in colorectal cancer [33]. At first, we were worried that autophagy blockage would increase the expression of PD-L1 in colon cancer cells [14], which was a potential risk. However, our study showed that JPJDR, as a mixed preparation, not only did not increase the expression of PD-L1 due to autophagy blockage, but also reduced the expression of PD-L1 in HT-29 cells. Since the anti-PD-1 therapy failed in clinical trials of colorectal cancer, and the model we used was a non-immune mouse, we cannot determine the therapeutic effect of JPJDR downregulating PD-L1 expression in tumor cells in vivo, which is what we will study next.

## 5. Conclusion

In summary, we confirmed the efficacy of Jianpi Jiedu Recipe in treating colon cancer in vitro and in vivo, and revealed the mitochondrial dysfunction and incomplete autophagy induced by it. At the same time, this study identified a rare formula that blocks autophagy in colon cancer while reducing its PD-L1 expression. These findings will provide more evidence and guidance for the clinical application of JDJDR.

## Abbreviations

JPJDR: Jianpi Jiedu Recipe
CRC: colorectal cancer
TCM: traditional Chinese medicine
ROS: reactive oxygen species
CQ: chloroquine

## Author contribution statement

Conception and design: Lingling Cheng, Jianlin Xu

Acquisition of data (provided animals, acquired and provided facilities): Lingling Cheng, Liangfeng Xu, HuaYuan, Qihao Zhao, Wei Yue

Analysis and interpretation of data: Shuang Ma, Xiaojing Wu, Dandan Gu, Yurong Sun, Haifeng Shi

Writing and review the manuscript: Lingling Cheng

Study supervision: Jianlin Xu

All data were generated in-house, and no paper mill was used. All authors agree to be accountable for all aspects of work ensuring integrity and accuracy

## Acknowledgement

Thank you for the financial support from “2023 Medical Scientific Research Project of Yancheng Health Commission (YK2023074)” and the equipment support from Yancheng Traditional Chinese Medicine Hospital.

